# Multimodal Imaging Mass Spectrometry of Murine Gastrointestinal Tract with Retained Luminal Content Shows Molecular Localization Patterns

**DOI:** 10.1101/2021.10.03.462819

**Authors:** Emma R. Guiberson, Aaron G. Wexler, Christopher J. Good, Eric P. Skaar, Jeffrey M. Spraggins, Richard M. Caprioli

## Abstract

Digestive diseases impact 62 million people a year in the United States. Despite the central role of the gut to human health, past imaging mass spectrometry (IMS) investigations into the gastrointestinal tract are incomplete. The gastrointestinal tract, including luminal content, harbors a complex mixture of microorganisms, host dietary content, and immune factors. Existing imaging approaches remove luminal content, and images focus on small regions of tissue. Here, we demonstrate the use of a workflow to collect multimodal imaging data for both intestinal tissue and luminal content. This workflow for matrix-assisted laser desorption/ionization imaging mass spectrometry retains luminal content and expands the amount of tissue imaged on one slide. Results comparing tissue and luminal content show unique molecular distributions using multimodal imaging modalities including protein, lipid, and elemental imaging. Leveraging this method to investigate intestinal tissue infected with *Clostridioides difficile* compared to control tissue shows clear differences in lipid abundance of various lipid classes in luminal content during infection. These data highlight the potential for this approach to detect unique biological and markers of infection in the gut.

## INTRODUCTION

Within the United States, over 62 million people are diagnosed with a digestive disease annually.^1^ The human gastrointestinal (GI) tract is the largest interface between the host, gut microbiota, and environmental factors within the body, which plays a major role in human health and disease.^2^ The GI tract consists of the small intestine and large intestine, with the small intestine including the duodenum, jejunum and ileum. All of these regions have unique functions and molecular characteristics, owing in part to the dramatically different bacterial burden in proximal and distal regions of the intestine.^3^ In addition to tissue, the gastrointestinal tract includes luminal content which contains bacteria from the gut microbiota, dietary molecules such as ingested fats, as well as pathogenic bacteria such as *Clostridioides difficile*.^4,5^ Matrix assisted laser desorption/ionization (MALDI) imaging mass spectrometry (IMS) has been employed to provide spatially-associated chemical information for molecular species in mammalian GI tissue.^6–8^ These studies do not encompass the entirety of the GI tract, as either an incomplete portion of the tissue is probed or the luminal content is removed. Sample preparation of this organ must be improved to leverage MALDI IMS for analysis of all regions of the GI tract. When preparing tissue for cryosectioning, murine intestines can be manipulated into a “Swiss roll” confirmation in which intestinal tissue is rolled into a spiral pattern.^9,10^ This approach enables imaging of more intestinal tissue (∼16 cm of total 33 cm in length) and allows comparisons between additional regions of the GI tract. Unfortunately, traditional “Swiss roll” protocols remove luminal content.^10^ Feces have a high fat and water composition, which leads to difficulties with slide retention during sample preparation protocols, like aqueous tissue washes, for MALDI IMS.^11^ Thus, we present an intestinal sample workflow for multimodal MALDI IMS of the murine GI tract, which includes the luminal content, to reintroduce the full complexity of the gut for deeper molecular coverage.

## EXPERIMENTAL

Instrument and sample preparation specifics included in supplemental Tables 1 and 2.

### Sample Preparation

Intestines were rolled into a spiral shape and embedded in 2.6% carboxymethylcellulose (CMC). Tissue and CMC were snap frozen over liquid nitrogen at −80°C, cryosectioned at 10 μm thickness using a CM3050 S cryostat (Leica Biosystems, Wetzlar, Germany) and thaw-mounted onto 1% poly-l-lysine coated ITO slides (Delta Technologies, Loveland, CO) for lipid and protein imaging, or vinyl slides for elemental imaging (VWR, Radnor, PA). Autofluorescence microscopy images were acquired using EGFP, DAPI and DsRed filters on a Zeiss AxioScan Z1 slide scanner (Carl Zeiss Microscopy GmbH, Oberkochen, Germany) prior to matrix application.

### MALDI FT-ICR Protein Imaging

Samples for protein analysis were washed using graded ethanol washes (70% EtOH, 100% EtOH, Carnoy’s Fluid, 100% EtOH, H_2_O, 100% EtOH) and Carnoy’s Fluid (6 ethanol: 3 chloroform: 1 acetic acid) to remove salts and lipids. Samples were sprayed with 2′,6′-Dihydroxyacetophenone (DHA). Following matrix application, high-mass-resolution imaging mass spectrometry of whole intestinal samples were performed using a Solarix 15T MALDI FT-ICR mass spectrometer (Bruker Daltonics, Billerica, MA). Data were visualized using SCiLS Lab 2020 (Bruker Daltonics, Billerica, MA).

### LA-ICP Elemental Imaging

Trace element imaging was performed as previously described.^12^ Samples were ablated using an LSX-213 laser ablation system (Teledyne CETAC, Omaha, NE, USA) and analyzed using a coupled Element 2 high-resolution sector field ICP-MS (Thermo Fisher Scientific, Waltham, MA) operated in medium-resolution mode. Helium gas was used to assist in transport of ablated sample particles from the ablation chamber to the ICP-MS. The resulting data were converted into vender-neutral imzML format and visualized using SCiLS Lab 2020.

### MALDI timsTOF Lipid Imaging

All samples for lipid analysis were washed with ammonium formate and distilled H_2_O prior to matrix applicated. 1,5-diaminonapthalene (DAN) matrix was sublimated onto the tissue. Imaging mass spectrometry was performed using a prototype timsTOF Pro equipped with a dual ESI/MALDI and operated in qTOF mode with TIMS deactivated.^13^ All tentative lipid identifications were determined based on mass accuracy using the LIPIDMAPS lipidomics gateway (lipidmaps.org) with a 5 ppm mass tolerance.

### MALDI FTICR Lipid Imagin

Samples were sprayed with 1,5-diaminonapthalene (DAN). Following matrix application, high-mass-resolution imaging mass spectrometry of intestinal samples were performed using a Solarix 15T MALDI FTICR mass spectrometer. A CASI method was used with a Q1 mass of *m/z* 790.00 and an isolation window of *m/z* 325 to reduce abundant taurocholate peak at *m/z* 514.29.

## RESULTS AND DISCUSSION

### Imaging of proximal and distal regions of GI tissue with intact luminal content

Regions of the GI tract can have dramatically different molecular makeups, due in part to the higher microbial burden in the distal intestine.^3^ Microbiota contribute greatly to the molecular diversity within the GI tract, including unique metabolites and interactions with dietary fats. The length of the GI tract prevents analysis of the entire tissue on one slide. The small intestine of a mouse is approximately 33 cm long, greatly surpassing the length on a traditional glass slide (typically 75 mm).^14^ The Swiss roll conformation, as shown in Figure 1, allows for half (∼16 cm) of this organ to be imaged at once with only one section. The section shown in Figure 1 illustrates this method with intact luminal content as used in our imaging workflow.

**Figure 1:**
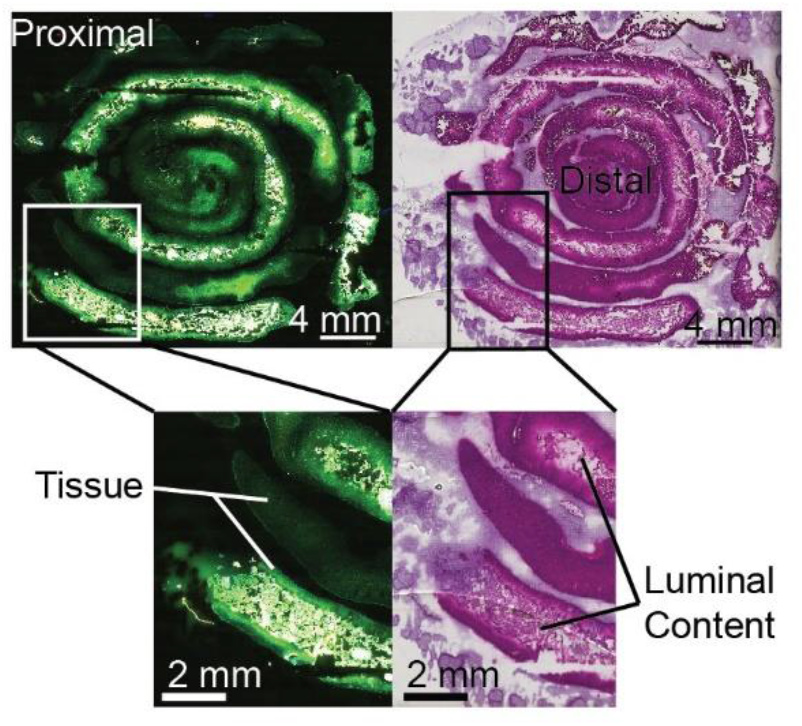
Autofluorescence and hematoxylin & eosin stains of adapted Swiss roll conformation. Microscopy images of a proximal intestine sample with intact luminal content in a Swiss roll conformation.

Luminal content adds additional sample preparation considerations for imaging mass spectrometry analysis. Especially with washing protocols, there is loss of luminal content and/or poor tissue retention with glass and ITO-coated slides. To address this sample loss, a polylysine coating was applied to indium tin oxide (ITO) slides.^15^ ITO slides, with and without a polylysine coating, and glass microscope slides were compared for tissue and luminal content retention following two different washing protocols (Figure S1). For lipid imaging, retention was tested after an aqueous ammonium formate (AF) wash to remove salt adducts (Figure S1a). For protein imaging, retention was tested following a Carnoy wash, consisting of graded ethanol washes and Carnoy’s fluid to remove lipids (Figure S1b).^16^ On a glass slide there was some tissue dropout, luminal content was removed with an AF wash, and luminal content was lost during a Carnoy wash. An ITO surface did not provide enough adherence to maintain luminal content during both washes. While a small amount of luminal content was lost with a polylysine coating and a Carnoy wash, retention greatly improved with an AF wash. Overall, polylysine improved tissue and luminal content retention after an ammonium formate wash and to a lesser degree with a Carnoy wash. These data suggests that utilizing a polylysine coating when working with intestinal tissue increases the spatial integrity of the sample.

### Tissue and luminal content in the gastrointestinal tract show unique multimodal molecular profiles

To fully elucidate the molecular differences between gut tissue and luminal content, a variety of IMS modalities were applied to proximal intestinal tissues. Using the polylysine coating, and respective washing methodologies, unique distributions of proteins, lipids, and elements were identified which alludes to the molecular complexity of the gut microenvironment. Autofluorescence images of these tissues are shown in Figure S2. Figure 2a shows proteins that localize either exclusively to the tissue (top row) or to luminal content (bottom row), highlighting the unique molecular profile of the luminal content and its distinction from host-only samples. Figure 2b shows various phosphatidylcholines with distinct localization patterns, despite being from the same lipid class, suggesting a difference between host lipids and diet-derived fecal lipids. Figure 2c shows elemental imaging contrasting dietary metals in the feces (calcium, manganese) against elements within the tissue (phosphorus), or elements present in both regions (magnesium, copper). These multimodal IMS images highlight the reproducibility of polylysine coating for tissue retention, as well as the value in studying luminal content alongside intestinal tissue as a unique microenvironment previously understudied. The addition of luminal content offers deeper understanding of the gastrointestinal tract and how it changes as a result of various conditions such as diet and disease.

**Figure 2:**
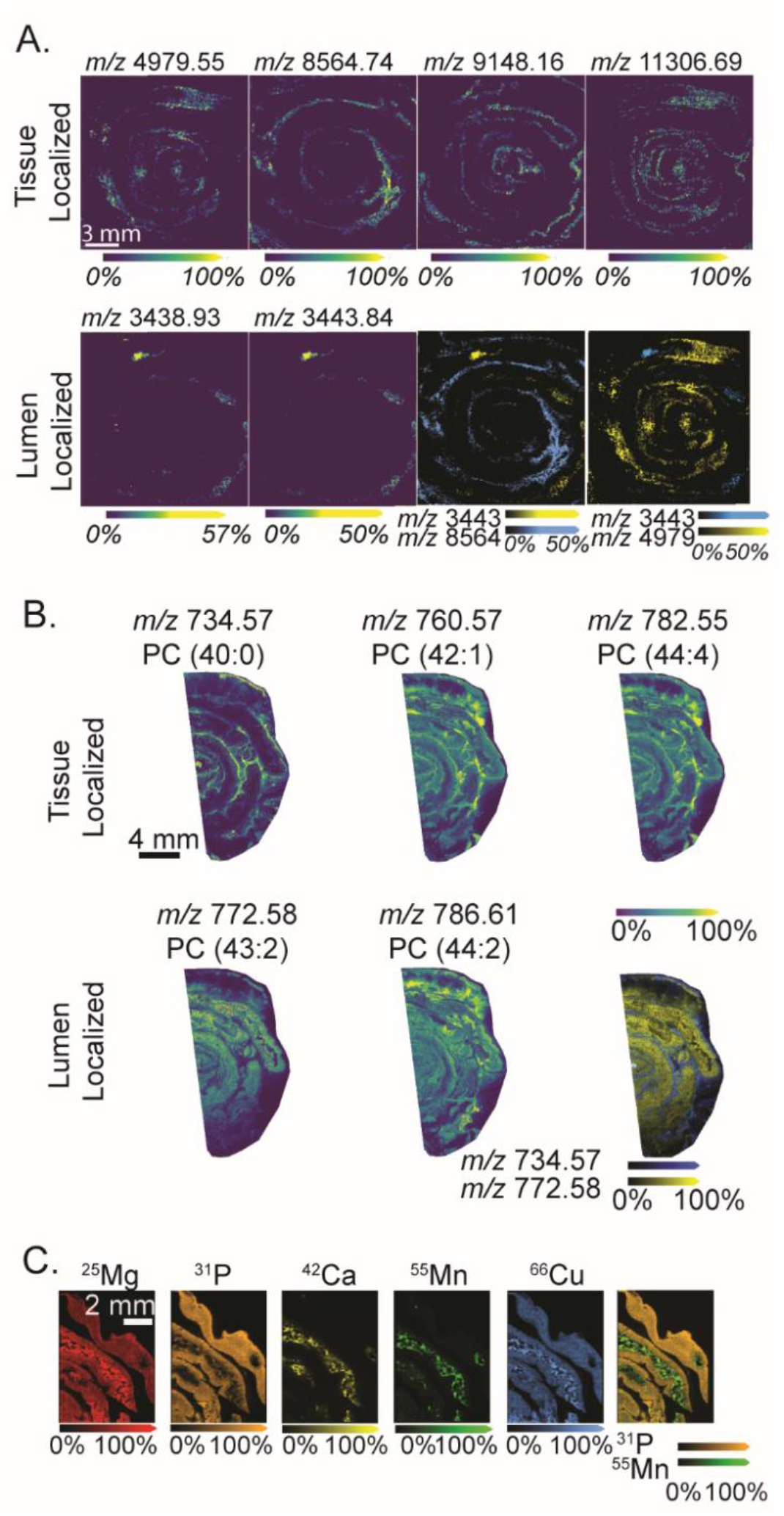
Multimodal IMS shows unique molecular profiles between tissue and luminal content regions. A) Protein imaging shows tissue-specific localized proteins (top row) and luminal content-specific proteins (bottom row) and overlaid proteins. B) Elemental imaging using LA-ICP IMS shows unique elemental distributions between tissue-specific (P), lumen-specific (Ca, Mn), or present in both regions (Mg, Cu) and overlaid elements. C) Lipid imaging showing tissue-specific localization (left) or lumen-specific localization (right) and overlaid lipids.

### A case study: fecal lipids differ in *C. difficile* infection model

To demonstrate the potential for luminal content in detecting novel molecular differences during disease, lipid abundances between *Clostridioides difficile-*infected and uninfected proximal intestinal tissue were compared. *C. difficile* spores localize to luminal content, then germinate into vegetative cells that cause inflammation through toxin production.^4^ Due to the impact of *C. difficile* on the gut, and spore localization patterns, this was an ideal infection model for these assays. Figure 3 shows clear differences in three lipids (PE(30:0) at *m/z* 662.47, PI(34:0) at *m/z* 837.54, IPC(36:0) at *m/z* 824.56) that are abundant in control samples but below the limit of detection in infected tissue. Due to their lumen localization, it is likely that these lipids are diet-derived, suggesting *C. difficile* infection impacts the breakdown of lipids in the gut. Additional use of this method has also shown a dramatic increase in taurocholate (*m/z* 514.29) abundance in luminal content during *C. difficile* infection,^17^ which would have gone undetected with a traditional approach. This case study demonstrates the potential applications of this multimodal workflow for biomarker analysis of fecal content, as shown by the clear molecular changes in luminal content during gastrointestinal disease.

**Figure 3:**
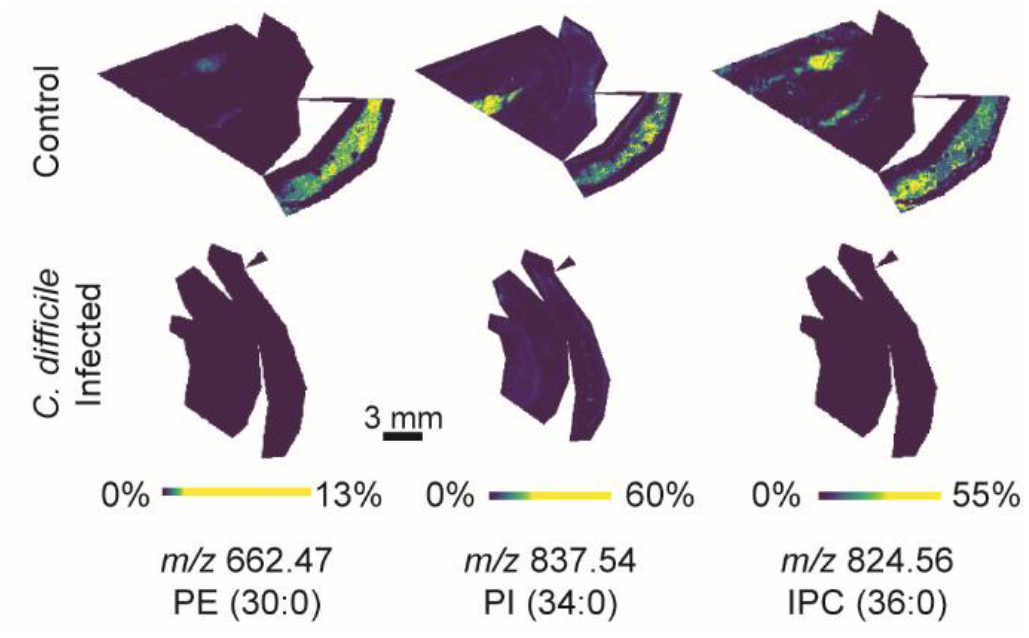
Dietary lipid profiles in luminal content change during *C. difficile* infection. MALDI IMS of control proximal intestinal samples with clear lumen localization of lipids (top) and same lipids in infected tissue (bottom).

Overall, this workflow to collect multimodal imaging data from intestinal tissue with luminal content offers a more comprehensive insight into the gastrointestinal tract through improved tissue coverage and luminal content retention. This application can be applied to further our understanding of various digestive diseases, and how the complex dynamic of the GI tract may change from the internal and external factors that act upon it.

## SUPPORTING INFORMATION

Additional experimental details and methods, including autofluorescence images of main text figures tissue and washing experiments.

## ACKNOWLEDGMENTS AND FUNDING SOURCES

We gratefully acknowledge lab members for review of this manuscript, and Dr. William Perry for his assistance with elemental imaging. This work was supported by National Institute of Allergy and Infectious Diseases grant R01AI45992 and R01AI145992 (to. E.P.S. and J.M.S.). The prototype MALDI timsTOF MS was developed as part of the National Science Foundation Major Research Instrument Program (CBET – 1828299 awarded to J.M.S. and R.M.C.) and the Bruker 15T solariX FT-ICR MS in the Mass Spectrometry Research Center at Vanderbilt University was acquired through the NIH Shared Instrumentation Grant Program (1S10OD012359 awarded to R.M.C.). A.G.W. was supported by a fellowship from the Helen Hay Whitney Foundation.

## REFERENCES

1. Peery, A. F. et al. Burden and Cost of Gastrointestinal, Liver, and Pancreatic Diseases in the United States: Update 2018. Gastroenterology 156, 254-272.e11 (2019).

2. Thursby, E. & Juge, N. Introduction to the human gut microbiota. Biochem. J. 474, 1823–1836 (2017).

3. Hillman, E. T., Lu, H., Yao, T. & Nakatsu, C. H. Microbial ecology along the gastrointestinal tract. Microbes Environ. 32, 300–313 (2017).

4. Rineh, A., Kelso, M. J., Vatansever, F., Tegos, G. P. & Hamblin, M. R. Clostridium difficile infection: molecular pathogenesis and novel therapeutics. Expert Rev Anti Infect Ther 12, 131–150 (2015).

5. Li, Z. et al. Effects of metabolites derived from gut microbiota and hosts on pathogens. Front. Cell. Infect. Microbiol. 8, (2018).

6. Kaya, I., Jennische, E., Lange, S. & Malmberg, P. Analytical Methods Dual polarity MALDI imaging mass spectrometry on the same pixel points reveals spatial lipid localizations at high-spatial resolutions in rat small intestine †. Anal. Methods 10, 2428–2435 (2018).

7. Genangeli, M. et al. MALDI-Mass Spectrometry Imaging to Investigate Lipid and Bile Acid Modifications Caused by Lentil Extract Used as a Potential Hypocholesterolemic Treatment. J. Am. Soc. Mass Spectrom. (2019). doi:10.1007/s13361-019-02265-9

8. Carter, C. L. et al. Characterizing the Natural History of Acute Radiation Syndrome of the Gastrointestinal Tract: Combining high mass and spatial resolution using MALDI-FTICR-MSI Claire. Health Phys. 116, 454–472 (2020).

9. Bialkowska, A. B., Ghaleb, A. M., Nandan, M. O. & Yang, V. W. Improved swiss-rolling technique for intestinal tissue preparation for immunohistochemical and immunofluorescent analyses. J. Vis. Exp. 2016, 1–8 (2016).

10. Pereira, A. et al. Comparison of two techniques for a comprehensive gut histopathological analysis : Swiss Roll versus Intestine Strips. Exp. Mol. Pathol. 111, 104302 (2019).

11. Jensen, R., Buffangeix, D. & Covi, G. Measuring water content of feces by the Karl Fischer method. Clin. Chem. 22, 1351–1354 (1976).

12. Becker, J. S. et al. Bioimaging of metals by laser ablation inductively coupled plasma mass spectrometry (LA-ICP-MS). Mass Spectrom. Rev. 29, 156–175 (2010).

13. Spraggins, M. et al. High-Performance Molecular Imaging with MALDI Trapped Ion-Mobility Time-of-Flight (timsTOF) Mass Spectrometry. (2019). doi:10.1021/acs.analchem.9b03612

14. Hugenholtz, F. & Vos, W. M. De. Mouse models for human intestinal microbiota research : a critical evaluation. Cell. Mol. Life Sci. 75, 149–160 (2018).

15. Mazia, D., Schatten, G. & Sale, W. Adhesion of Cells to Surfaces Coated with Polylysine. J. Cell Biol. 66, 198–200 (1975).

16. Yang, J. & Caprioli, R. M. Matrix Sublimation/Recrystallization for Imaging Proteins by Mass Spectrometry at High Spatial Resolution. Anal. Chem. 83, 5728–5734 (2011).

17. Wexler, A. G. et al. Clostridioides difficile infection induces a rapid influx of bile acids into the gut during colonization of the host ll Clostridioides difficile infection induces a rapid influx of bile acids into the gut during colonization of the host. CellReports 36, 109683 (2021).

