## Supplemental Figures and Tables for "Multimodal Imaging Mass Spectrometry of Murine Gastrointestinal Tract with Retained Luminal Content Shows Molecular Localization Patterns"


**AUTHOR LINE**

Emma R. Guiberson[1,2], Aaron G. Wexler[3,4], Christopher J. Good[1,2], Eric P. Skaar[3,4], Jeffrey M. Spraggins[1,2,5,6]*, Richard M. Caprioli[1,2,5,7,8]

**AUTHOR AFFILIATIONS**

1 Mass Spectrometry Research Center, Vanderbilt University, Nashville, TN, USA, 37203
2 Department of Chemistry, Vanderbilt University, Nashville, TN, USA, 37203
3 Vanderbilt Institute for Infection, Immunology, and Inflammation, Vanderbilt University Medical Center, Nashville, TN, USA, 37203
4 Department of Pathology, Microbiology, and Immunology, Vanderbilt University Medical Center, Nashville, TN, USA, 37203
5 Department of Biochemistry, Vanderbilt University, Nashville, TN, USA, 37203
6 Department of Cell and Developmental Biology, Vanderbilt University, Nashville, TN, USA, 37203
7 Department of Medicine, Vanderbilt University, Nashville, TN, USA, 37203g
8 Department of Pharmacology, Vanderbilt University, Nashville, TN, USA, 37203

*Corresponding author




**KEYWORDS:** MALDI IMS, Gastrointestinal tract

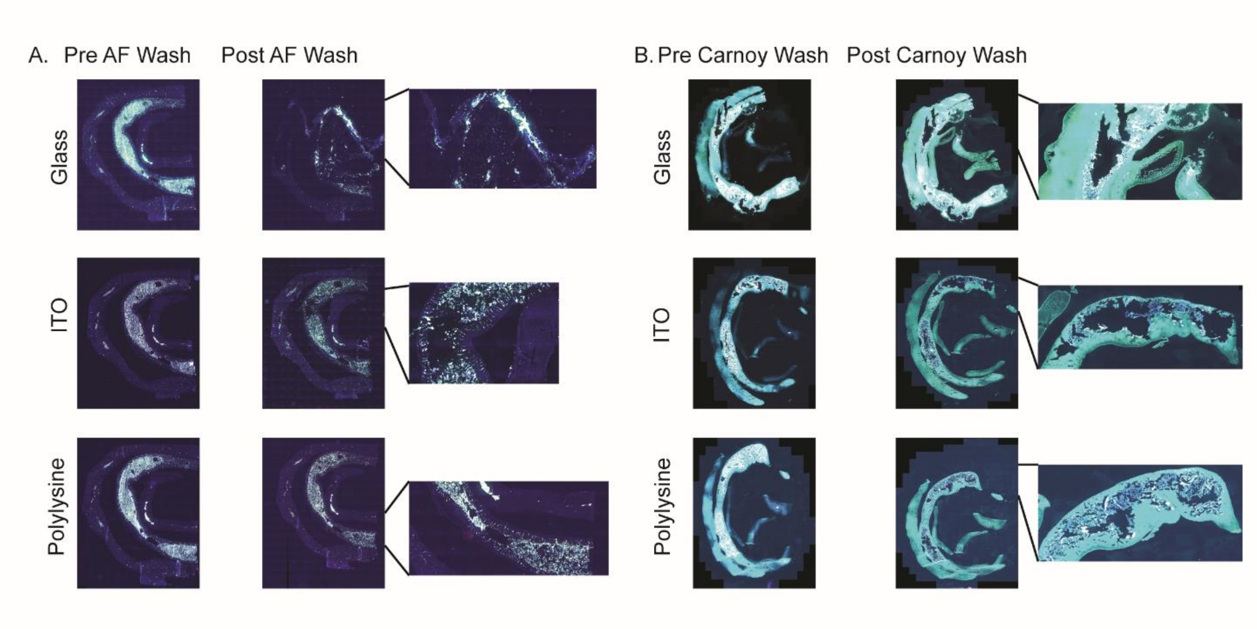


**Supplemental Figure 1: Poly-l-lysine coating improved luminal content retention for various washing methods.** A) Pre- and post-washing autofluorescence images on glass, ITO and polylysine-coated ITO slides using an ammonium formate wash. B) Pre- and post-washing autofluorescence images in glass, ITO and polylysine-coated ITO slides using a Carnoy wash.

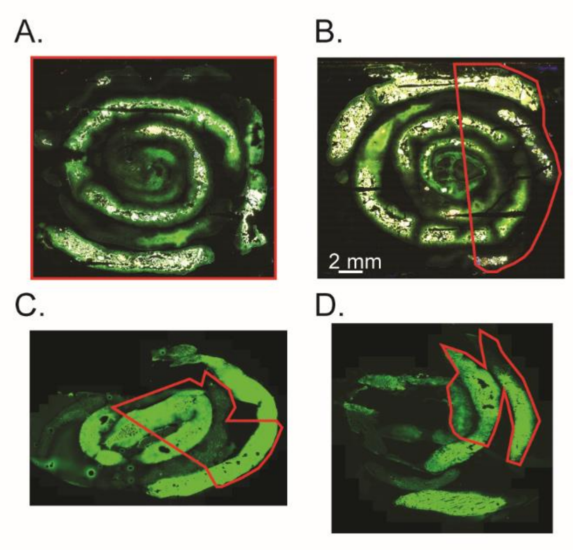


**Supplemental Figure 2: Autofluorescence images of imaged tissue showing regions ablated.** A) Tissue sample used for protein imaging in Figure 2A, B) Tissue sample used for lipid imaging in Figure 2C, C) Control tissue sample used for lipid imaging in Figure 3, D) Infected tissue sample used for lipid imaging in Figure 3.

**Supplemental Table 1**

| | MALDI FTICR Protein Imaging | LA-ICP IMS Elemental Imaging | MALDI timsTOF Lipid Imaging | MALDI FTICR Lipid Imaging |
|---|---|---|---|---|
| **Laser** | Apollo II dual MALDI/ESI ion source and a Smartbeam II 2 kHz frequency tripled Nd:YAG laser 355 nm[18] | Nd:YAG 213 nm | Nd:YAG 355 nm | Nd:YAG 355 nm |
| **Laser Setting** | small (50 um) | 100 um | small (50 um) | large (~150 um) |
| **Spatial Resolution** | 100 um | 60 um | 25 um | 75 um |
| **Laser Shots** | 1000 | - | 400 | 300 |
| **Laser Power** | 70% | 35% | 40% | 35% |
| ***m/z* range** | 1000-30,000 | - | 200-2000 | 300-2000 |
| **Resolving Power** | 36,507 at *m/z* 8529.91899 | - | - | 71,928 at *m/z* 885.5498 |
| **Transient length** | 2.3069s | - | - | 1.251s |
| **Ionization mode** | positive | - | positive | negative |

**Supplemental Table 2**

| | **MALDI FTICR Protein Imaging** | **MALDI timsTOF Lipid Imaging** | **MALDI FTICR Lipid Imaging** |
|---|---|---|---|
| **Matrix** | 2′,6′-Dihydroxyacetophenone (DHA) | 1,5-diaminonapthalene (DAN)[19] | 1,5-diaminonapthalene (DAN) |
| **Matrix Solution** | 90 mg/mL DHA in 70% ACN with 100 µL TFA and 50 µL of 30% ammonium hydroxide | 300 mg DAN (~1.0 mg/cm$^2$) | 20 mg/mL DAN in THF |
| **Application Method** | Sprayed with automatic robotic aerosol sprayer (TM Sprayer, HTX Technologies, Chapel Hill, NC, USA) | Sublimated using a simple sublimation apparatus at 130°C and 24 mTorr for 3 min[20] | Sprayed with automatic robotic aerosol sprayer (TM Sprayer, HTX Technologies, Chapel Hill, NC, USA) |
| **TM Sprayer Method** | 45°C, 1050 mm/min, 0.075 mL/min, 1.5 mm track spacing, 8 passes | -- | 40°C, 1100 mm/min, 0.05 mL/min, 1.5 mm track spacing, 5 passes, 75°C heated stage |

**SUPPLEMENTAL METHODS:**

**Materials:** Acetic acid, 1,5-diaminonapthalene (DAN), trifluoroacetic acid (TFA), ammonium formate, 2′,6′-Dihydroxyacetophenone (DHA), hematoxylin, and eosin were purchased from Sigma-Aldrich Chemical Co. (St. Louis, MO, USA). HPLC-grade acetonitrile, methanol, ethanol, and chloroform were purchased from Fisher Scientific (Pittsburgh, PA, USA).

**Histology.** Following MALDI IMS experiments, matrix was removed from samples using 100% ethanol and rehydrated with graded ethanol and double distilled $H_2O$. Tissue samples were stained using a hematoxylin and eosin stain, and brightfield microscopy of stained tissues was obtained using a Leica SCN400 Brightfield Slide Scanner at 20x magnification (Leica Microsystems, Buffalo Grove, IL).

**Animal Protocols:** All animal experiments were performed using protocols approved by the Vanderbilt University Medical Center Institutional Animal Care and Use Committee. Animal studies were conducted using 8- to 12-week old male C57Bl/6 (Jackson Laboratories) mice. Mice infected with *C. difficile* were given cefoperazone in their drinking water (0.5 mg/ml) for 5 days. Following a 2-day recovery period, mice were gavaged orally with $10^5$ *C. difficile* spores or PBS to an endpoint of three days.

**Supplemental References:**


18. Prentice, B. M., Chumbley, C. W. & Caprioli, R. M. High-speed MALDI MS/MS imaging mass spectrometry using continuous raster sampling. *J. Mass Spectrom.* **50**, 703–710 (2015).
19. Thomas, A., Charbonneau, J. L., Fournaise, E. & Chaurand, P. Sublimation of new matrix candidates for high spatial resolution imaging mass spectrometry of lipids: Enhanced information in both positive and negative polarities after 1,5-diaminonapthalene deposition. *Anal. Chem.* **84**, 2048–2054 (2012).
20. Hankin, J. A., Barkley, R. M. & Murphy, R. C. Sublimation as a Method of Matrix Application for Mass Spectrometric Imaging. *J Am Soc Mass Spectrom* **18**, 1646–1652 (2007).
